# Thermal cycling-hyperthermia down-regulates liposaccharide-induced inflammation in human blood and helps with clearance of herpes simplex virus type 1

**DOI:** 10.1101/2022.12.20.521318

**Authors:** Yu-Yi Kuo, Guan-Bo Lin, Wei-Ting Chen, You-Ming Chen, Hsu-Hsiang Liu, Chih-Yu Chao

## Abstract

Infection would lead to temperature increase in the affected region or entire human body, in order to weaken the pathogens, such as virus, or activate the immune system. As an alternative therapy with extensive application for various diseases, hyperthermia (HT) can regulate the release of pro-inflammatory cytokines and the antiviral activity of immune system. However, existing studies have found that overheating impairs healthy tissues and immune cells. The study puts forth a modified HT treatment, thermal cycling-hyperthermia (TC-HT), looking into its effect on immunomodulation and cellular viabilities. It shows that TC-HT can reduce the secretion of pro-inflammatory cytokines, induced by lipopolysaccharide (LPS) both *ex vivo* and *in vitro*, and elevate the efficacy of U-937 macrophages in clearing herpes simplex virus type 1 (HSV-1) *in vitro*. Furthermore, via optimizing its parameters, TC-HT can boost the efficacy of U-937 macrophage in clearing HSV-1, which may be attributed to the enhancement of actin polymerization and phagocytosis activity via TC-HT. In sum, TC-HT outperforms HT in safety and therapeutic effect in immunomodulation, shedding light on its potential in the treatment of immunological diseases.

## Introduction

As ubiquitous pathogens, viruses are the cause of many diseases, in both animals and humans, including lethal and devastating ones, such as influenza [1], acquired immunodeficiency syndrome (AIDS) [2], Ebola [3], and coronavirus disease 2019 (COVID-19) [4]. There have been some treatments for virus-induced diseases, such as ribavirin and oseltamivir, which regulate viral activities by interfering with the replication process [5] and restricting replicated viruses in cytoplasm [6], respectively. Besides, certain viral diseases, such as smallpox [7] and hepatitis B [8], can be prevented by vaccination, which trains the adaptive immune system to produce specific immunoglobulin against the viruses. Whatsoever, the host immune system still plays the vital role in combating infection.

It is a common knowledge that the innate immune system is the first-line defense against invading microbial pathogens, such as viruses. The innate immunity first identifies pathogens via receptors on the surface of innate immune cells, such as macrophages, dendritic cells, and neutrophils [9]. The activated immune cells as well as the infected cells release chemokines and cytokines as chemical messengers. Chemokines are molecules producing chemotaxis to recruit immune cells to the infected site, while cytokines modulate the innate immune responses by regulating the differentiation, proliferation and activation of immune cells or uninfected cells [10]. For example, tumor necrosis factor-alpha (TNF-α), a common cytokine after infection, can activate nuclear factor kappa-light-chain-enhancer of activated B cells (NF-κB) and mitogen-activated protein kinase (MAPK) pathways, which are related to cell differentiation, proliferation and apoptosis [11]. Interferon (IFN), a cytokine secreted by infected cells, can warn uninfected cells and inhibit viral replication [12]. Besides, there is a subclass of cytokines interleukin (IL) mediating the interactions between leukocytes and playing an important role in bridging the innate and adaptive immune responses [13]. Cytokines would be secreted after infection becomes pro-inflammatory and induces inflammation, which increases blood flow and vascular permeability. Then, leukocytes and plasma proteins infiltrate the infected sites, causing typical inflammatory symptoms, such as topical redness, heat, and swelling [14]. Meanwhile, inflammation also triggers phagocytosis, an important innate immune response to clear pathogens. Infiltrated macrophages and dendritic cells at the infected sites would devour the pathogens and break them down into molecular patterns, which are then presented to lymphocytes such as T cells, activating adaptive immune response and hence generating host immunity [15].

Hyperthermia (HT), a conventional alternative therapy, boasts extensive clinical applications for various diseases due to its pleiotropic effect on mammalian cells. For instance, existing studies have proven that it can inhibit cancer cell proliferation and induce apoptosis, in addition to synergistically amplifying the anticancer effect of some chemotherapy drugs and phytochemical molecules [16, 17]. Besides, proper heat stress can activate heat shock proteins and antioxidant pathways, producing neuroprotective effects [18], and heat is conducive to immunity. It has also been proven that enhancement of core body temperature can lower the mortality related to several viral infections [19]. Moreover, heat can up-regulate immune function, such as the phagocytic potential of macrophages and dendritic cells [20, 21], while boosting adaptive immune response [19]. However, HT could also harm health tissues, especially those with lower thermal tolerance. While skin cells, for instance, can tolerate up to 47 °C in temperature [22], central and peripheral nervous systems in both human [23, 24] and animal models [25, 26] can be damaged by temperature over 43 °C. Prolonged heat exposure can suppress the innate immunity of mouse model by decreasing the number of macrophages and retarding the maturation of dendritic cells [27]. Overall, continuing application of conventional HT can be detrimental to normal healthy tissues and cells, necessitating its modification in clinical application.

Our previous study put forth a new heat treatment method, named thermal cycling-hyperthermia (TC-HT), with a process consisting of a desired high-temperature stage and a low-temperature stage alternately. Administered along with anticancer agents, TC-HT exhibited a stronger inhibitive effect on pancreatic cancer cells than HT, via modulating cell death pathways, such as apoptosis and cell cycle arrest [28-30]. Meanwhile, the treatment doesn’t affect normal cells. Our study also showed that compared with HT, TC-HT can provide better neuroprotection against the cytotoxicity of both hydrogen peroxide and beta-amyloid, via regulating the ROS level and continually activating the neuroprotective pathways, without damaging neural cells as conventional HT often does [31]. Therefore, as promising substitute for HT, TC-HT could provide stronger and safer immunomodulatory effect than continuous HT.

In this paper, the study applied TC-HT treatment to both the *ex vivo* model (human blood) and the *in vitro* model (U-937 macrophages) with lipopolysaccharide (LPS), and the *in vitro* model of U-937 macrophages with herpes simplex virus type 1 (HSV-1), looking into its immunomodulatory effect and comparing it with HT. It finds that, in the *ex vivo* model, TC-HT reduced the LPS-induced secretion of pro-inflammatory cytokines TNF-α, interleukin-6 (IL-6), and interleukin-1 beta (IL-1β) at a level higher than HT in both whole blood and isolated peripheral blood mononuclear cells (PBMCs), while better preserving the cell viability of PBMCs than HT. In the *in vitro* model, TC-HT also outperforms HT in preserving the cell viability of U-937 macrophages and activating U-937 macrophages for HSV-1 clearance, the latter of which can be further augmented by optimizing the efficacy of U-937 macrophages for clearing HSV-1, via altering the low-temperature duration of TC-HT treatment to 2 min. The optimized TC-HT treatment demonstrated a greater efficacy on inhibiting LPS-induced TNF-α and IL-6 secretions, while promoting actin polymerization levels and phagocytosis activity in U-937 macrophages. In summary, TC-HT has the potential to activate more efficient immune responses, and provide less damage and discomfort to patients.

## Materials and Methods

### HT and TC-HT applications

All HT and TC-HT treatments were administered by a modified polymerase chain reaction (PCR) machine (Applied Biosystems; Thermo Fisher Scientific, Inc., Waltham, MA, USA), as shown in Fig 1A. For HT treatment, the high temperature setting can be chosen to be 39, 42, or 45 °C applied continuously for 30 min. The TC-HT treatment was composed by a 10-cycle repeated procedure of a desired high-temperature stage for a period of time and followed by a low-temperature stage. In this study, the setting temperature of the high-temperature stage was set at 42 or 45 °C for 3 min, and the low-temperature stage was set at 37 °C for a certain duration (Fig 1B). The actual temperature of both HT and TC-HT treatments was measured via a type-K thermocouple and a temperature/process controller (PZ900, RKC instrument Inc., Tokyo, Japan). The thermocouple was located at the bottom of the cell culture well, and the monitored actual temperature was recorded every 10 sec, as shown in Fig 1C.

**Fig 1.**
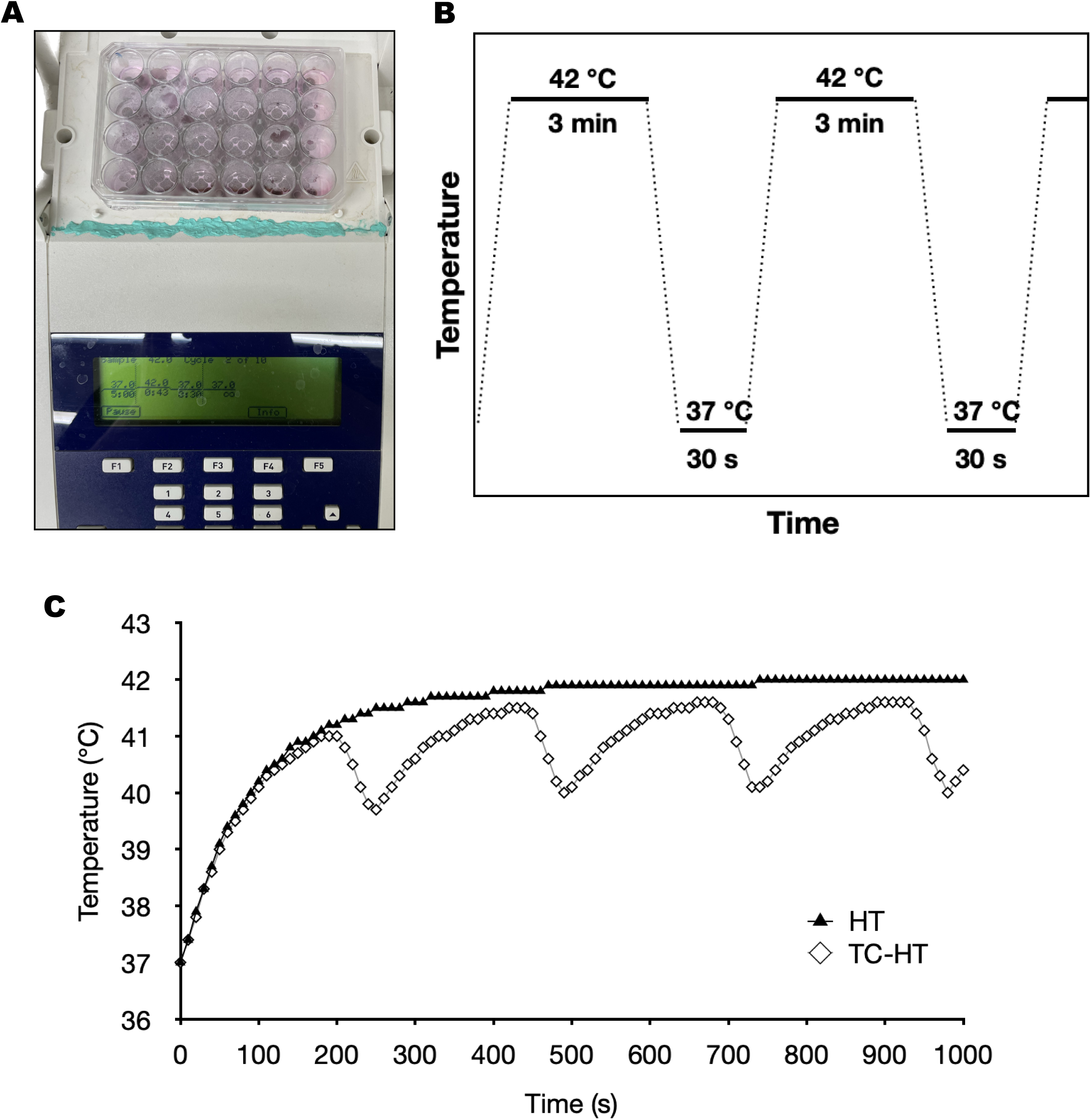
Experimental setups for thermal treatments. (A) TC-HT and HT treatments were applied by a PCR machine. (B) The schematic representation of the setting parameters of TC-HT and (C) the actual temperatures measured every 10 sec during thermal treatments by the thermocouple.

### Whole blood collection

Venous blood samples were collected from the antecubital vein of six healthy volunteers using the BD Vacutainer^®^ blood collection tubes (cat. 367874, Becton Dickinson, Franklin Lakes, NJ, USA) containing 158 IU of sodium heparin. Informed consent was obtained from all the volunteers prior to the experiments. All volunteers did not receive any medication (antibiotics or anti-inflammatory agents) or any therapy. For whole blood assay, 1:3 blood dilutions were made with sterile Roswell Park Memorial Institute (RPMI) 1640 medium (HyClone; GE Healthcare Life Sciences, Chicago, IL, USA) supplemented with 100 unit/mL penicillin and 100 mg/mL streptomycin (Gibco Life Technologies, Grand Island, NY, USA).

### PBMCs isolation

The PBMCs in blood were isolated by Ficoll-1077 density gradient centrifugation. 1:1 blood dilution was made with sterile phosphate buffered saline (PBS) (HyClone). The dilutions were slowly layered onto Ficoll-1077 (Sigma-Aldrich; Merck KGaA, Darmstadt, Germany) and centrifuged at 400×g for 30 min at room temperature without brake. After centrifugation, the PBMCs layer between the plasma and Ficoll was withdrawn carefully, and then rinsed and maintained with sterile RPMI-1640 medium supplemented with 100 unit/mL penicillin and 100 mg/mL streptomycin. The cell density was counted and the PBMCs were cultured in a final concentration of 1.5×10^6^ cells/mL in 24- or 96-well culture plates (Thermo Fisher Scientific, Inc., Waltham, MA, USA) for the following experiments.

### Thermal treatments and inflammation by LPS

To investigate the immunomodulatory effects of the HT and TC-HT treatments on blood, PBMCs, and U-937 macrophages, 50 ng/mL LPS (Sigma-Aldrich; Merck KGaA) was used to stimulate inflammatory response. For the post-treatment group, LPS was treated 1 h prior to the HT or TC-HT treatment. For the pre-treatment group, LPS was treated 4 h after the HT or TC-HT treatment. Samples without LPS treatment were used as the control group. At 6 h after the LPS treatment, the blood or PBMCs samples of each group were centrifuged at 2000×g for 5 min. The supernatants were collected and stored at -20 °C until the enzyme-linked immunosorbent assay (ELISA) experiment.

### ELISA experiment

Sandwich ELISA experiments for LPS-induced TNF-α, IL-6, and IL-1β were carried out using ELISA MAX^TM^ Standard Set (Biolegend, San Diego, CA, USA). In brief, the detected cytokine was sandwiched by the corresponding capture antibody and detection antibody in accordance with the manufacturer’s instructions in the microwell plates for ELISA (Corning Inc., Corning, NY, USA). After the targeted inflammatory cytokines were sandwiched, 100 μL TMB substrate (Biolegend) was used for color development in dark for 15-30 min. The color development was stopped by 100 μL sulfuric acid (Echo Chemical Co. Ltd., Miaoli, Taiwan) of concentration 1 M. The absorbance at 450 nm was measured by Multiskan GO microplate spectrophotometer (Thermo Fisher Scientific, Inc.), and the reference absorbance at 690 nm was subtracted. The concentration of the inflammatory cytokines was calculated based on the standard calibration curve, which was independently determined in each assay according to manufacturer’s instructions.

### Cell and virus culture

Human monocytic cell line U-937 and African green monkey kidney epithelial cell line Vero were purchased from Biosource Collection and Research Center (Hsinchu, Taiwan) and cultured at 37 °C in a humidified incubator with 5% CO_2_. U-937 cells were maintained in RPMI-1640 medium supplemented with 4.5 g/L glucose, 10 mM 4-(2-hydroxyethyl)-1-piperazineethanesulfonic acid (HEPES, 1.0 mM sodium pyruvate (Sigma-Aldrich; Merck KGaA), 10% fetal bovine serum (Hyclone) and 1% of penicillin-streptomycin (Gibco). Vero cells were maintained in Minimal Essential Medium with Earle’s Balanced Salts Solution (MEM/EBSS) (Hyclone), supplemented with 1.0 mM sodium pyruvate (Sigma-Aldrich; Merck KGaA), 10% fetal bovine serum (Hyclone), 0.1 mM non-essential amino acids and 1% penicillin-streptomycin (Gibco). The HSV-1 strain KOS (ATCC VR-1493, American Type Culture Collection, Manassas, VA, USA) was propagated in Vero cells, and was aliquoted and stored at -80 °C.

### Monocytes differentiation into macrophages

For differentiation, U-937 cells (2×10^5^ cells/mL) were incubated in the culture medium containing 200 ng/mL of phorbol myristate acetate (PMA) (Sigma-Aldrich; Merck KGaA) for 24 h to induce differentiation into macrophage cells. After the PMA treatment, the adherent U-937 macrophages were rinsed with PBS and resuspended in medium. Centrifugation at 200×g for 8 min was performed for cell collection. After counting the cell density, U-937 macrophages were cultured in 24-well culture plates (5×10^5^ cells/well) and incubated at 37 °C in a humidified incubator with 5% CO_2_ for 48 h prior to the HT and TC-HT treatments.

### MTT assay

The viabilities of the PBMCs or U-937 macrophages were analyzed by 3-(4,5-dimethylthiazol-2-yl)-2,5-diphenyltetrazolium bromide (MTT) (Sigma-Aldrich; Merck KGaA) assay. The MTT powder was dissolved in distilled water at a concentration of 5 mg/mL as the stock solution. At 24 h after the HT or TC-HT treatment, cells were incubated in 100 µL of their culture medium with a final MTT concentration 0.5 mg/mL at 37 °C for 4 h. Cells without HT or TC-HT treatment were used as the untreated control group. Formazan was crystalized after the MTT incubation, and 100 µL solubilizing buffer of 10% sodium dodecyl sulfate (SDS) (Bioshop Canada Inc., Burlington, ON, Canada) solution in 0.01 N hydrochloric acid (HCl) (Echo Chemical Co. Ltd.) was used to dissolve the formazan crystals at 37 °C overnight in the dark. As the crystals in each well were fully dissolved, the absorbance of each well was detected by Multiskan GO microplate spectrophotometer (Thermo Fisher Scientific, Inc.). The quantity of formazan of each well was determined by the absorbance at 570 nm, and the reference absorbance at 690 nm was subtracted. The cell viabilities were expressed in percentage and the untreated control was set at 100%.

### Plaque reduction assay and U-937 macrophage infection

The viabilities of HSV-1 after heating or infecting U-937 macrophage were detected by plaque reduction assay. Vero cells for plaque reduction assay were cultured in 24-well culture plates (1.5×10^5^ cells/well) for 24 h before the plaque reduction assay. To analyze the plaque formation ability, Vero cells were incubated with the heated HSV-1 or cell lysates of HSV-1-infected macrophages at 37 °C for 1h. After incubation, medium contained HSV-1 was removed and the infected Vero cells were overlaid with the Vero culture medium containing 0.5% methylcellulose (Sigma-Aldrich; Merck KGaA) for 4 days. After the incubation, Vero cells were fixed with 4% paraformaldehyde (PFA) (Sigma-Aldrich; Merck KGaA) for 10 min and stained with 0.1% crystal violet (Sigma-Aldrich; Merck KGaA). To verify the effect of thermal treatment on the formation of HSV-1 plaques in Vero cells, the plaque numbers formed by the heated or non-heated HSV-1 were first compared. Fig 2 showed that the plaque formation in Vero cells was significantly reduced as HSV-1 was heated by 42 °C HT for 30 min before infection, suggesting that heat exposure could inhibit HSV-1 viability. Therefore, to exclude the influence of heating on the formation of HSV-1 plaques in Vero cells, thermal treatments were applied to U-937 macrophages before the HSV-1 infection. Furthermore, to assess the effect of HT and TC-HT treatments on the viral clearance ability, U-937 macrophages were infected with HSV-1 at 0.1 multiplicity of infection (MOI) 1 h after the HT and TC-HT treatments. The infected U-937 macrophages without any thermal treatment were used as the control group. At 24 h after the HSV-1 infection, the culture medium of U-937 macrophages was collected, and cells were then lysed in the culture medium of Vero cells for three freeze-thaw cycles. Cell lysates were collected and used in plaque reduction assay. Total numbers of plaques in each well of each group were counted and compared to the control group, whose plaque formation percentage was set at 100%.

**Fig 2.**
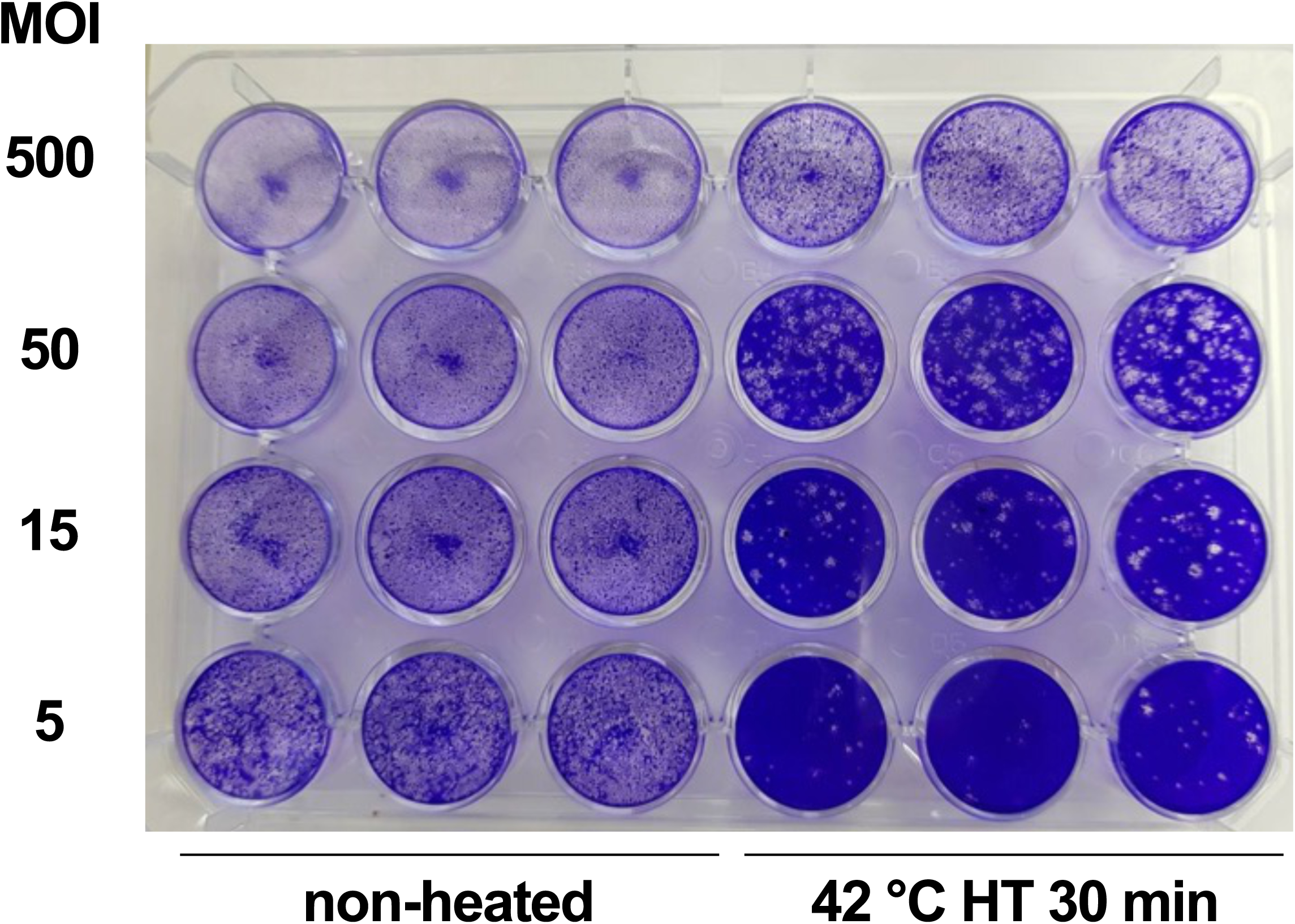
Effect of TC-HT and HT treatments on HSV-1 infectivity *in vitro*. Effects of thermal treatment on HSV-1 infectivity in Vero cells were assessed by plaque reduction assay. MOI: multiple of infection.

### Immunofluorescence staining and laser scanning confocal microscopy

To observe the effect of rearranging filamentous-actin (F-actin) by HT and TC-HT treatments on U-937 macrophages, Phalloidin-iFluro 488 reagent (ab176753, Abcam, Cambridge, MA, USA) was used for labeling F-actin according to manufacturer’s instruction. In brief, 5×10^5^ U-937 cells were grown on coverslips for thermal treatments. Then, the cells were rinsed three times with PBS and fixed with 4% PFA at 24h after thermal treatments. After fixation, cells were permeabilized with 0.1% Triton X-100 (Bioshop Canada Inc.) for 5 min and incubated with 1% bovine serum albumin (BSA) (Bioshop Canada Inc.) in PBS for 30 min to block non-specific sites for improving staining. Cells were stained with Phalloidin-iFluro 488 Reagent (1:1000 in 1% BSA in PBS) for 1 h at room temperature in dark, and were rinsed with PBS for three times before mounting the coverslips to slides in glycerol mounting medium with 4’,6-diamidino-2-phenylindole (DAPI) (ab188804, Abcam). The mounted samples were observed on a Zeiss LSM 880 inverted laser scanning confocal microscope. The Phalloidin-iFluro 488 (green fluorescence) and DAPI (blue fluorescence) staining dyes were excited at a wavelength of 488 nm and 405 nm, respectively. Image analysis was performed using ImageJ v3.91 software for calculating the relative fluorescence units (RFU) of green and blue fluorescence. Actin polymerization is defined as the normalized RFU ratio of green/blue fluorescence (iFluro 488/DAPI) relative to the control group [32].

### Phagocytosis activity assay

FITC-dextran (MW 40,000, Sigma-Aldrich; Merck KGaA) was used to assess the effect of HT and TC-HT treatments on phagocytosis activity of U-937 macrophages. A stock of 20 mg/mL FITC-dextran was diluted in PBS, and a reaction concentration of 1 mg/mL was incubated with the U-937 macrophages at 24 h after HT and TC-HT treatments at 37 °C in a humidified incubator with 5% CO_2_ for 1h. Untreated macrophages were used as the positive control, incubated with FITC-dextran at 37 °C in a humidified incubator with 5% CO2. Negative control samples were incubated at 4 °C in the dark to assess non-specific binding of U-937 macrophages to FITC-dextran. Fluorescence signals were measured using FACSCanto II system (BD Biosciences, San Jose, CA, USA) with the FL1 channel. The mean fluorescence level of the negative control was subtracted from each group to exclude non-specific binding, and then the phagocytosis activity of FITC-dextran in U-937 macrophages was determined by normalizing to the positive control group.

### Statistical analysis

The results were presented as the mean ± standard deviation. Statistical analysis using one-way analysis of variance (ANOVA) followed by Tukey’s post-hoc test was performed by OriginPro 2015 software (OriginLab Corporation, Northampton, MA, USA). Results were considered to be statistically significant when P-values were less than 0.05.

## Results

### TC-HT post-treatment down-regulates the LPS-induced inflammatory response *ex vivo*

In the study, the regulatory effects of HT post-treatment on the LPS-induced pro-inflammatory cytokines in blood were first assessed. As shown in Figs 3A-C analyzed via ELISA assay, LPS treatment was found to greatly elevate the concentrations of TNF-α (from 47.76 ± 1.02 to 3390.61 ± 445.53 pg/mL), IL-6 (from 143.72 ± 8.42 to 3998.96 ± 312.03 pg/mL), and IL-1β (from 8.42 ± 0.02 to 89.24 ± 2.46 pg/mL) in whole blood. The elevations were further boosted up by 39 °C HT post-treatment (TNF-α from 3390.61 ± 445.53 to 4296.10 ± 158.27 pg/mL, IL-6 from 3998.96 ± 312.03 to 4103.02 ± 122.04 pg/mL, and IL-1β from 89.24 ± 2.46 to 212.9 ± 1.17 pg/mL, respectively), and the secretions of TNF-α, IL-6 and IL-1β stimulated by LPS were greatly inhibited by 45 °C HT post-treatment (TNF-α from 3390.61 ± 445.53 to 233.42 ± 7.59 pg/mL, IL-6 from 3998.96 ± 312.03 to 131.94 ± 2.14 pg/mL, and IL-1β from 89.24 ± 2.46 to 9.10 ± 0.04 pg/mL, respectively). On the other hand, 42 °C HT post-treatment showed down-regulatory effects on TNF-α (from 3390.61 ± 445.53 to 2498.09 ± 593.01 pg/mL) and IL-6 (from 3998.96 ± 312.03 to 3378.4 ± 92.2 pg/mL), but not on IL-1β (from 89.24 ± 2.46 to 140.33 ± 2.06 pg/mL). However, it was noted that the releases of all the three pro-inflammatory cytokines were significantly decreased as the temperature of HT post-treatment increased, thus suggesting that higher temperature was more favorable for anti-inflammatory treatment.

**Fig 3.**
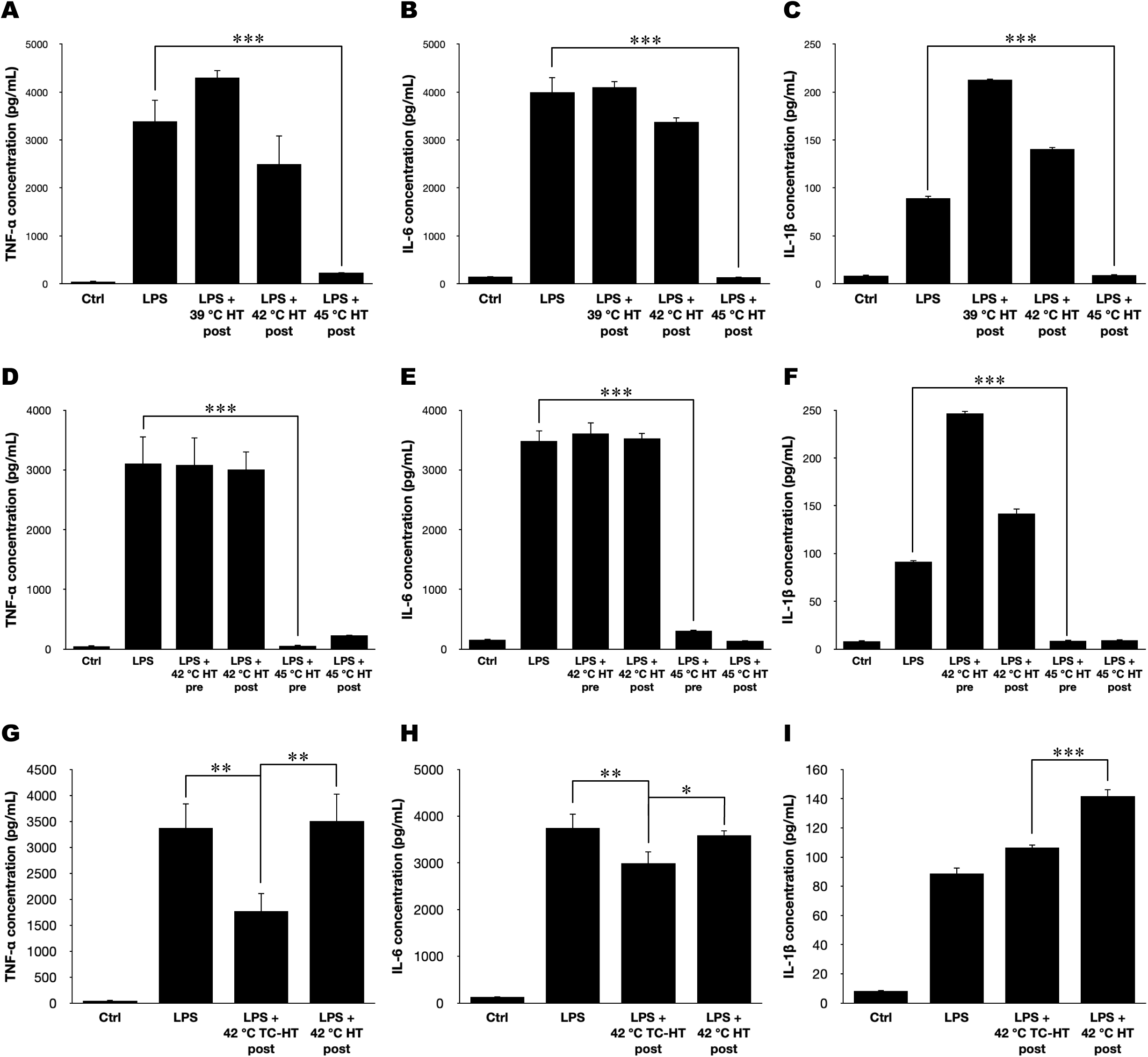
Effect of TC-HT and HT treatments on regulating the secretions of LPS-induced pro-inflammatory cytokines in whole blood *ex vivo*. The concentrations of LPS-induced TNF-α, IL-6, and IL-1β in whole blood were analyzed via ELISA assay in order to compare the immunomodulatory effects of (A-C) HT post-treatments, (D-F) HT pre-treatment and post-treatment, and (G-I) TC-HT and HT post-treatments. Data were presented as the mean ± standard deviation (n = 3 in each group). One-way ANOVA with Tukey’s post hoc test was used to determine statistical significance (*P < 0.05, **P < 0.01, and ***P < 0.001 between indicated groups). Pre: pre-treatment group; Post: post-treatment group.

Although 45 °C HT post-treatment performed strong inhibitory effects on LPS-induced inflammatory responses *ex vivo*, studies have also indicated that overheating could impair the viability of immune cells or their immunological functions [27], as a result of which the effect of high temperature on LPS-induced inflammatory response was further assessed by HT pre-treatment. Figs 3D-F showed that 42 °C HT pre-treatment did not alter LPS-induced secretion of TNF-α (3110.65 ± 450.34 and 3079.80 ± 466.87 pg/mL) and IL-6 (3486.53 ± 174.55 and 3610.29 ± 185.00 pg/mL), but promoting the secretion of IL-1β (from 91.52 ± 1.82 to 246.62 ± 2.72 pg/mL). This promotion by 42 °C HT pre-treatment on IL-1β was stronger than 42 °C HT post-treatment (246.62 ± 2.72 and 141.77 ± 5.13 pg/mL), but there was no significant difference on TNF-α (3079.80 ± 466.87 and 3007.42 ± 302.09 pg/mL) and IL-6 (3610.29 ± 185.00 and 3527.33 ± 89.09 pg/mL) between 42 °C HT pre-treatment and post-treatment. Noticeably, when blood was pre-treated by 45 °C HT, LPS was nearly unable to stimulate any inflammatory response in all the three cytokines, indicating that 45 °C HT treatment may dysregulate the inflammatory function of blood in response to LPS, and hence was not suitable for anti-inflammatory treatments. Thus, 42 °C was chosen as the setting temperature of the hyperthermic stage for the subsequent experiments.

In our previous studies, TC-HT showed higher therapeutic efficacies than HT for cancer treatment and neuroprotection [28, 30, 31]. As for the anti-inflammatory effect against LPS, 42 °C TC-HT post-treatment also showed a better performance than 42 °C HT post-treatment (Figs 3G-I). Secretions of TNF-α and IL-6 stimulated by LPS were significantly down-regulated by 42 °C TC-HT post-treatment (TNF-α from 3373.61 ± 474.96 to 1767.3 ± 350.90 pg/mL and IL-6 from 3748.51 ± 304.59 to 2999.39 ± 255.25 pg/mL), though IL-1β level was elevated slightly (from 88.74 ± 4.03 to 106.52 ± 2.05 pg/mL). Nevertheless, in LPS-induced TNF-α, IL-6 and IL-1β secretions, our results showed that 42 °C TC-HT post-treatment had significantly less inflammatory responses compared to 42 °C HT post-treatment, pointing out that TC-HT had more therapeutic potential than HT in anti-inflammatory treatment.

Inflammatory cytokines in blood were mainly secreted by immune cells, as a result of which we further isolated PBMCs to assess the regulatory effects of TC-HT and HT treatments on LPS-induced inflammatory response. As shown in Figs 4A-C analyzed via ELISA assay, 42 °C HT post-treatment was unable to inhibit LPS-induced PBMCs on releasing TNF-α (201.85 ± 10.29 and 239.57 ± 61.98 pg/mL) and IL-6 (305.00 ± 6.09 and 315.11 ± 10.63 pg/mL), but it had a down-regulation in IL-1β concentration (from 13.92 ± 0.26 to 10.10 ± 0.02 pg/mL). On the contrary, 42 °C TC-HT post-treatment on PBMCs significantly inhibited the secretion of TNF-α (from 201.85 ± 10.29 to 119.03 ± 3.50 pg/mL), IL-6 (from 305.00 ± 6.09 to 228.68 ± 4.49 pg/mL), and IL-1β (from 13.92 ± 0.26 to 10.30 ± 0.09 pg/mL) stimulated by LPS, while it caused a significant lower level than 42 °C HT post-treatment except for IL-1β in isolated PBMCs. Furthermore, we found that the viabilities of PBMCs at 24 h after 42 °C HT post-treatment were significantly lower than 42 °C TC-HT post-treatment (Fig 4D). Similar results were observed as the setting temperature of the hyperthermic stage was raised to 45 °C. Collectively, 42 °C TC-HT post-treatment was a safer and more effective anti-inflammatory treatment than 42 °C HT post-treatment against LPS *ex vivo*, and hence this treatment was chosen for the following immunomodulatory experiments.

**Fig 4.**
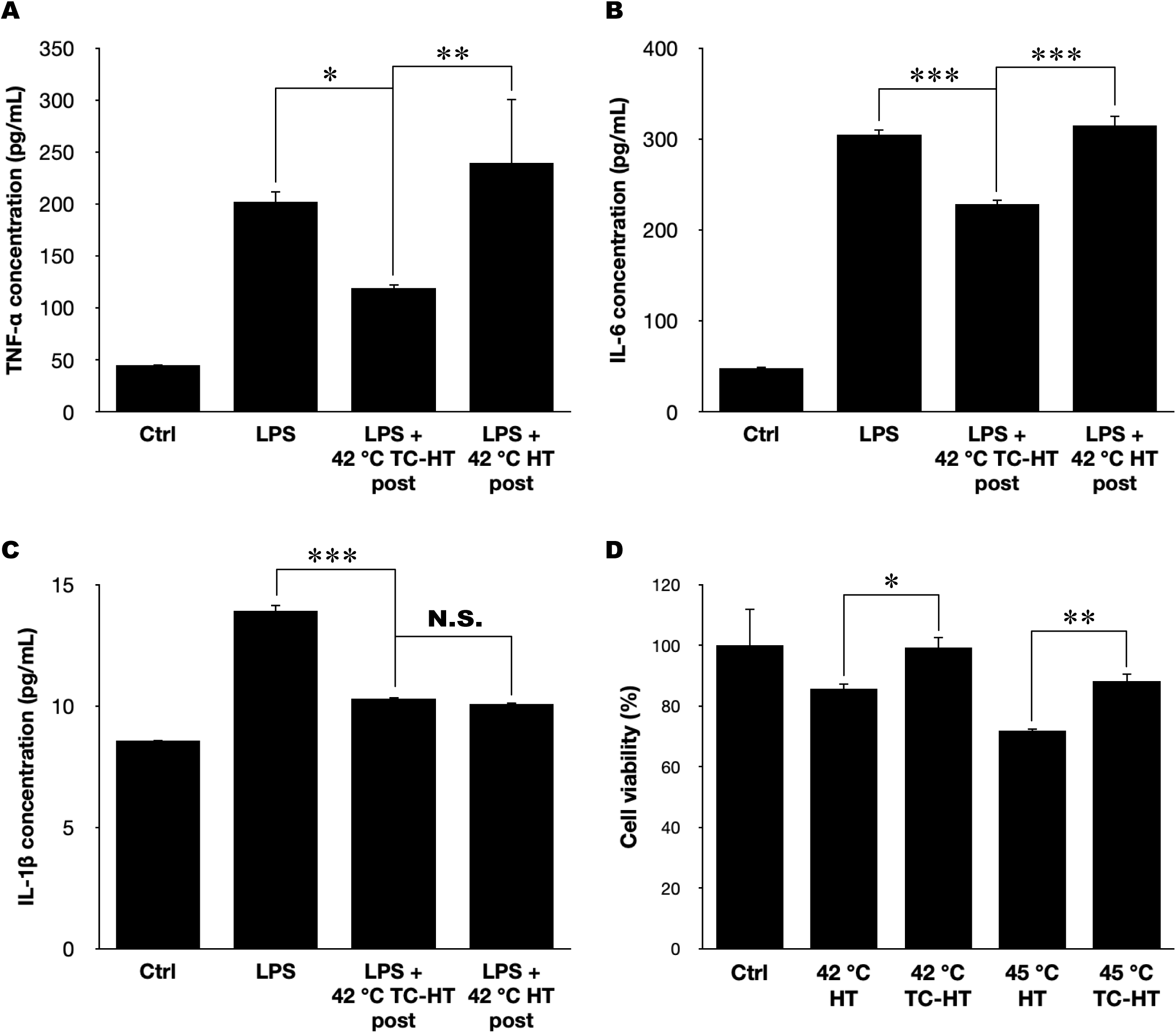
Effect of TC-HT and HT treatments on regulating the secretions of LPS-induced pro-inflammatory cytokines in PBMCs and on the cell viability of PBMCs *ex vivo*. (A-C) The concentrations of LPS-induced TNF-α, IL-6, and IL-1β in isolated PBMCs were analyzed via ELISA assay in order to compare the immunomodulatory effects of 42 °C TC-HT and HT post-treatments, and (D) the cell viability of PBMCs at 24 h after TC-HT and HT treatments was analyzed by MTT assay. Data were presented as the mean ± standard deviation (n = 3 in each group). One-way ANOVA with Tukey’s post hoc test was used to determine statistical significance (*P < 0.05, **P < 0.01, and ***P < 0.001 between indicated groups; N.S.: No significance between indicated groups). Post: post-treatment group.

### TC-HT treatment up-regulates macrophage’s viral clearance ability and down-regulates the LPS-induced inflammatory response *in vitro*

TC-HT has been shown to be capable of down-regulating LPS-induced inflammatory response *ex vivo*, and we then further investigated its regulatory effect on the viral clearance by U-937 macrophages *in vitro* and the anti-inflammatory effect against LPS stimulation in U-937 macrophages. The effects of TC-HT and HT treatments on U-937 macrophage viabilities were analyzed via MTT assay at 24 h after the thermal treatments. In Fig 5A, it was observed that 42 °C HT treatment decreased the viability of U-937 macrophages to 85.75%, while 42 °C TC-HT treatment preserved the viability at 100.43%. As the setting temperature of the hyperthermic stage was raised to 45 °C, HT treatment further reduced the viability of U-937 macrophages to 80.45%, and TC-HT treatment also reduced the viability to 82.94%. Therefore, TC-HT treatment was a safer thermal treatment than HT treatment, and 42 °C was chosen as the setting temperature of the hyperthermic stage for the following experiments.

**Fig 5.**
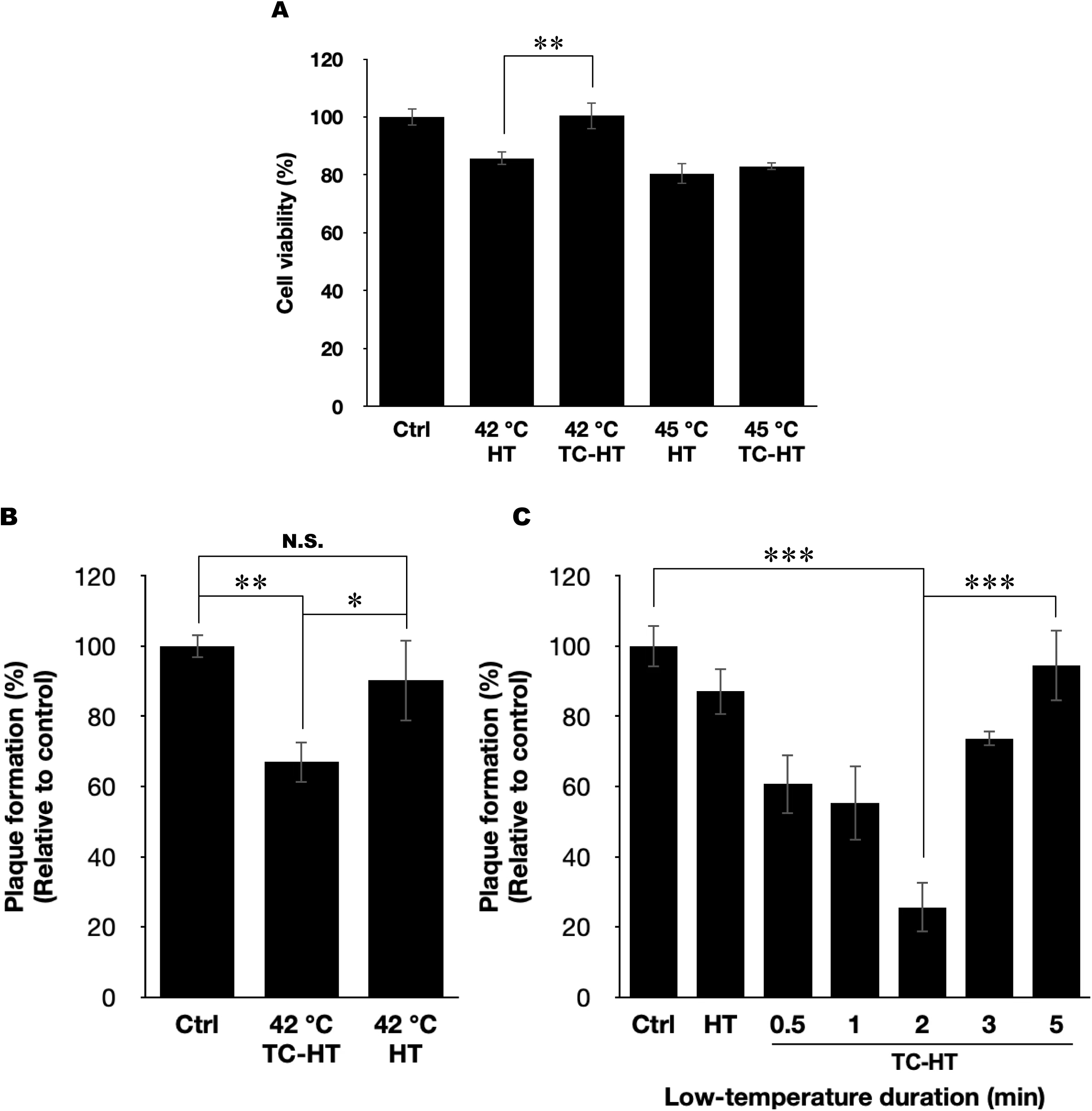
Effect of TC-HT and HT treatments on the viability of U-937 macrophages and HSV-1 clearance efficacy of U-937 macrophages *in vitro*. (A) Cell viabilities of U-937 macrophages at 24 h after TC-HT and HT treatments were analyzed by MTT assay. (B) Effects of thermal treatment on U-937 macrophages for clearing HSV-1 by HT and TC-HT and (C) by TC-HT with different low-temperature durations were assessed by plaque reduction assay. One-way ANOVA with Tukey’s post hoc test was used to determine statistical significance (*P < 0.05, **P < 0.01, and ***P < 0.001 between indicated groups; N.S.: No significance between indicated groups).

At 24 h after HSV-1 infection, the cell lysates of U-937 macrophages were collected as described in Materials and Methods section. The study found that the lysates of the TC-HT-treated U-937 macrophages formed significantly fewer plaques (66.99%) than that of the HT-treated U-937 macrophages (90.29%), as shown in Fig 5B. The result pointed out that 42 °C TC-HT treatment enhanced the efficacy of U-937 macrophages for clearing HSV-1 than 42 °C HT treatment *in vitro*. In addition to the importance of the setting temperature in the high-temperature stage of TC-HT, the duration of the low-temperature stage was also crucial for the neuroprotective effects in our previous study [31]. In this work, similar results were also observed in the plaque reduction assay, where the plaque formation was greatly inhibited down to 25.60% as the low-temperature duration in TC-HT extended to 2 min (Fig 5C), suggesting a stronger HSV-1 clearance efficacy by U-937 macrophages. However, this boost in the HSV-1 clearance diminished as the low-temperature duration was further extended beyond 3 min. These results indicated that 2 min was the optimal setting of low-temperature duration for 42 °C TC-HT treatment to clear HSV-1 by U-937 macrophages. For longer or shorter low-temperature durations, the efficacy for HSV-1 clearance by U-937 macrophages was unable to be fully established, therefore showing more HSV-1 plaques in Vero cells.

In addition, the anti-inflammatory effect of the optimized 42 °C TC-HT post-treatment was examined via ELISA assay. Similar to the results in the *ex vivo* model, LPS treatment greatly increased TNF-α (from 1113.68 ± 41.65 to 30524.20 ± 1901.18 pg/mL) and IL-6 (from 13160.56 ± 43.79 to 28609.56 ± 521.61 pg/mL) secretions in U-937 macrophages, as shown in Fig 6. The inflammatory responses were down-regulated by the optimized 42 °C TC-HT post-treatment, with significant decreases in both TNF-α (from 30524.20 ± 1901.18 to 10530.88 ± 1461.47 pg/mL) and IL-6 (from 28609.56 ± 521.61 to 24618.44 ± 1793.60 pg/mL). Importantly, the down-regulation of LPS-induced pro-inflammatory cytokines secretion by the optimized 42 °C TC-HT post-treatment was significantly stronger than 42 °C HT post-treatment on both TNF-α (10530.88 ± 1461.47 and 14523.12 ± 1226.48 pg/mL) and IL-6 (24618.44 ± 1793.60 and 28671.90 ± 414.21 pg/mL), suggesting that TC-HT exhibited greater anti-inflammation potential than HT in the *in vitro* U-937 macrophage model.

**Fig 6.**
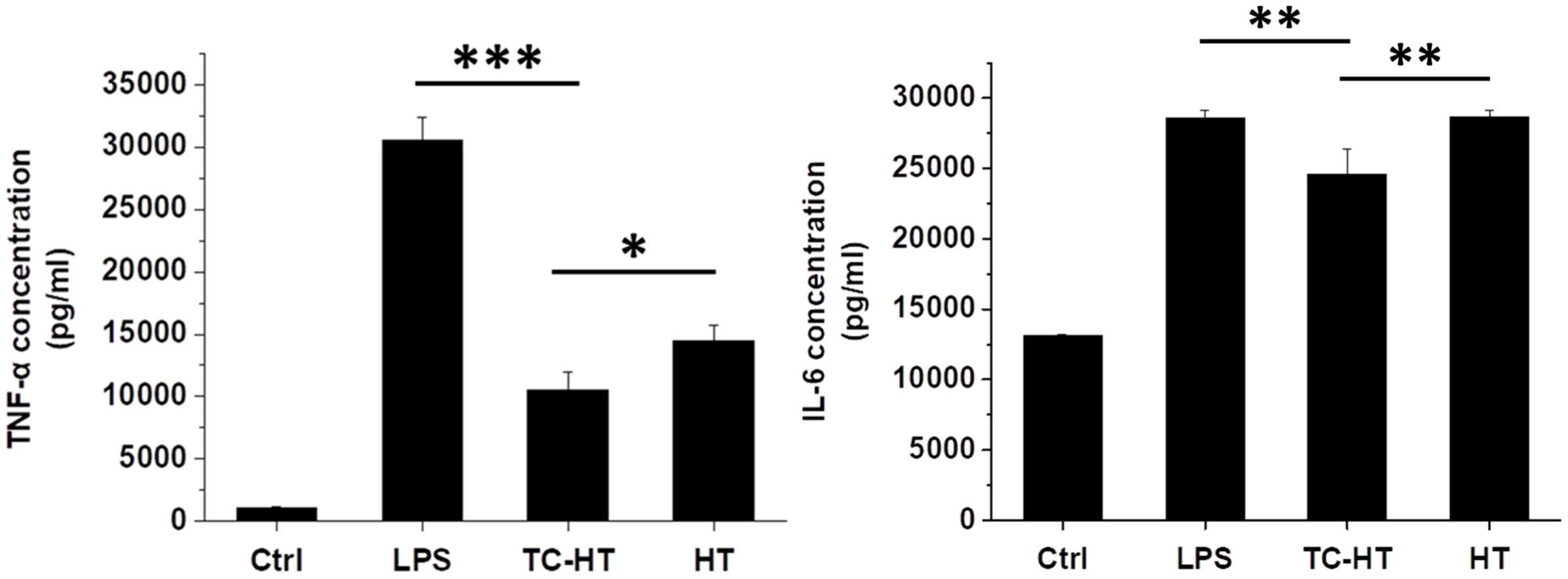
Effect of TC-HT and HT treatments on regulating the secretion of LPS-induced pro-inflammatory cytokines in U-937 macrophages. The concentrations of LPS-induced TNF-α and IL-6 in U-937 macrophages were analyzed via ELISA assay in order to compare the immunomodulatory effects of the optimized 42 °C TC-HT and HT post-treatments. Data were presented as the mean ± standard deviation (n = 3 in each group). One-way ANOVA with Tukey’s post hoc test was used to determine statistical significance (*P < 0.05, **P < 0.01, and ***P < 0.001 between indicated groups).

### TC-HT treatment elevates F-actin levels in U-937 macrophages

F-actin is the polymerized actin, and its level in cells is associated with the phagocytosis activity [33]. We used Phalloidin-iFluro 488 fluorescent dye to stain F-actin in the U-937 macrophages to analyze its relative levels, and the optimized 42°C TC-HT treatment with 2 min low-temperature duration (Fig 5C) was implemented on the cells. As shown in Fig 7, the RFU ratio of green/blue fluorescence was significantly higher in TC-HT-treated U-937 macrophages (1.37-fold increase compared to non-treated cells), indicating a significant elevation in actin polymerization by TC-HT treatment. Moreover, the result showed that the elevation of F-actin level in TC-HT-treated U-937 macrophages significantly outperformed that in HT-treated cells (0.92-fold decrease compared to the control cells). Combining with the viral clearance results in Fig 5, the study suggests that TC-HT may exert stronger phagocytosis activity via increasing actin polymerization in U-937 macrophage to clear HSV-1.

**Fig 7.**
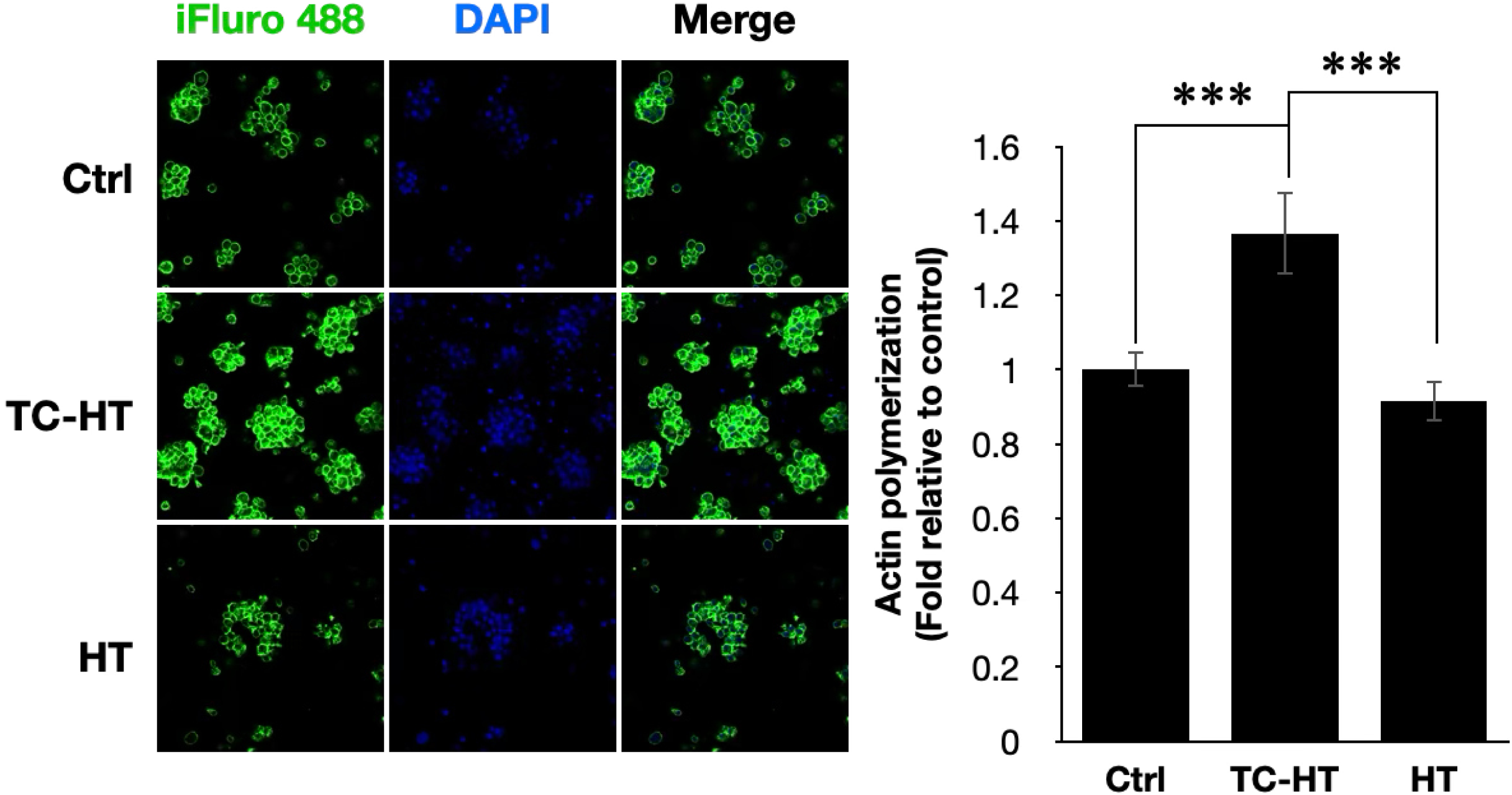
Effect of TC-HT and HT treatments on the level of F-actin in U-937 macrophages *in vitro*. F-actin and the nucleus of U-937 macrophages were stained respectively by Phalloidin-iFluro 488 fluorescent dye (green) and DAPI (blue) at 24 h after the TC-HT and HT treatments. Fluorescent signals were imaged by confocal microscope. Actin polymerization level of each group was determined by calculating the normalized RFU ratio of green/blue fluorescence (iFluro 488/DAPI) relative to the control group. Data were presented as the mean ± standard deviation (n = 4 in each group). One-way ANOVA with Tukey’s post hoc test was used to determine statistical significance (***P < 0.001 between indicated groups).

### TC-HT treatment promotes phagocytosis activity in U-937 macrophages

Macrophages and the immune cells play a crucial role in clearing pathogens, through phagocytosis, where they engulf and destroy pathogens including viruses [34]. Here, FITC-dextran was used as a surrogate pathogen to assess the phagocytosis activity of U-937 macrophages treated with TC-HT or HT. The results showed that, in comparison with the untreated control cells, a significantly promotion in phagocytosis activity (1.64-fold increase) was observed in U-937 macrophages treated with the optimized 42 °C TC-HT (Fig 8). Noteworthily, this promotion of phagocytosis activity significantly surpassed that by HT treatment (a 1.20-fold increase), indicating that the optimized 42 °C TC-HT treatment can enhance the defense of U-937 macrophages more effectively than HT treatment by increasing phagocytosis activity.

**Fig 8.**
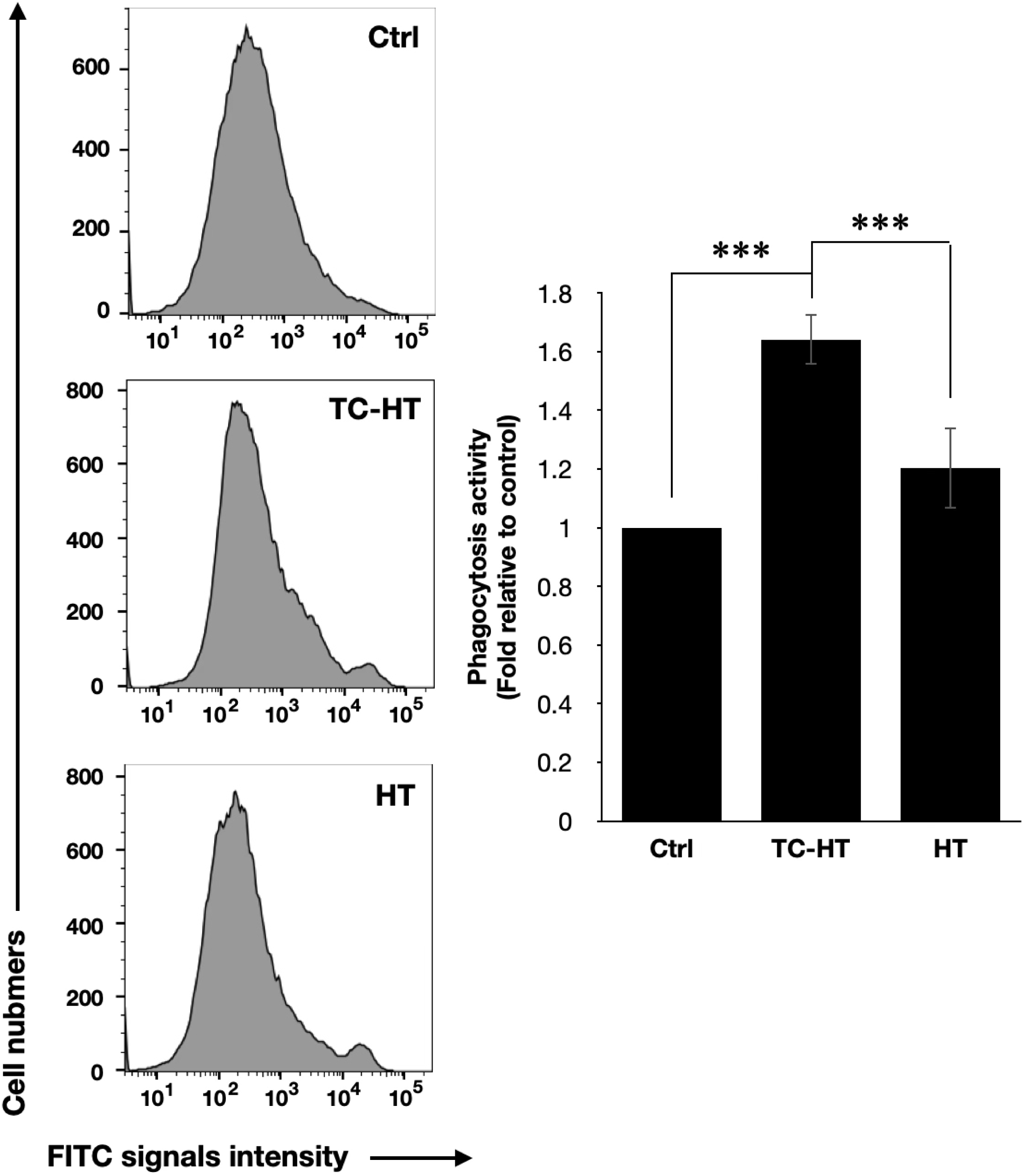
Effect of TC-HT and HT treatments on the phagocytosis activity in U-937 macrophages *in vitro*. Phagocytosis activity of U-937 macrophages at 24 h after the TC-HT and HT treatments was investigated with FITC-dextran. Fluorescent signals of the FITC-dextran in U-937 macrophages were detected using flow cytometry. Phagocytosis activity after each treatment was determined by the normalized mean fluorescence intensity of FL1 channel relative to the control group. Data were presented as the mean ± standard deviation (n = 4 in each group). One-way ANOVA with Tukey’s post hoc test was used to determine statistical significance (***P < 0.001 between indicated groups).

## Discussion

Inflammation is a common response in various diseases, a part of the immunological mechanism [35]. When immune cells identify exogenous pathogens, pro-inflammatory cytokines are secreted and more immune cells are summoned to the infected region. LPS, a large molecule on the outer membrane of Gram-negative bacteria [36], is a classical activator of inflammatory responses in monocytes and macrophages [37], prompting the cells to secrete pro-inflammatory cytokines such as TNF-α, IL-6, and IL-1β via NF-κB pathways [38]. TNF-α is an early mediator of inflammation, IL-1β is secreted from macrophages in infection [39, 40], while IL-6 is crucial to acute-phase response in inflammation [41]. As these cytokines are important indicators for inflammatory responses, the study employed them to assess inflammatory degrees.

As an alternative treatment with extensive applications in the treatment of cancer, cardiological diseases and physiotherapy [42], HT has extended its reach to immunomodulation, proven to be capable of inhibiting the release of pro-inflammatory cytokines in clinical study [43]. In our study, the results show that HT post-treatment with a setting temperature exceeding 42 °C decreased LPS-induced TNF-α, IL-6 and IL-1β levels in whole blood (Figs 3A-C). However, excessively high setting temperature (> 42 °C) of HT would cause dysfunctional inflammatory response to LPS (Figs 3D-F). In addition, both 42 and 45 °C HT post-treatments reduce the viability of isolated PBMCs significantly, which doesn’t occur in TC-HT post-treatments (Fig 4D), suggesting that TC-HT post-treatment is a better approach in immunomodulation of *ex vivo* system. Furthermore, 42 °C TC-HT post-treatment exhibits significant anti-inflammatory effects via suppressing LPS-induced TNF-α and IL-6 levels in both whole blood (Figs 3G and 3H) and isolated PBMCs (Figs 4A and 4B). For LPS-induced IL-1β, although the level is not inhibited, the elevation level of LPS-induced IL-1β after 42 °C TC-HT post-treatment is much less than corresponding 42 °C HT post-treatment (Fig 3I), while both suppress LPS-induced IL-1β in isolated PBMCs. These results manifest that TC-HT post-treatment is potentially a better anti-inflammatory therapy than HT post-treatment in *ex vivo*.

On top of regulating the inflammatory response against exogenous LPS, heat is also conducive to antiviral activities, as febrile temperature can not only reduce viral entry and replication [44] but also activate the innate and adaptive immune system, thus intensifying defensive activities [45]. As shown in Fig 2, after being treated with 42 °C HT for 30 min, HSV-1 directly generates less plaques in Vero cells than the non-heated HSV-1, suggesting a reduction on HSV-1 infectivity on Vero cells. On the other hand, the study finds that, after 42 °C TC-HT treatment, HSV-1 plaques formed by U-937 macrophage lysates are significantly less than that of 42 °C HT treatment (Fig 5B), indicating that 42 °C TC-HT treatment boosts U-937 macrophages’ efficacy in clearing HSV-1 *in vitro*. In addition, TC-HT also outperforms HT in upholding the viabilities of U-937 macrophages at setting temperatures 42 and 45 °C (Fig 5A). These results show that TC-HT is a safer and more effective treatment than HT in increasing the HSV-1 clearance efficacy for U-937 macrophages *in vitro*.

In TC-HT, low-temperature stages are interspersed into the continuous hyperthermic process, thereby outperforming HT in immunomodulatory effect both *ex vivo* and *in vitro*, perhaps due to more suitable low-temperature duration between high-temperature stages, which mitigates the adverse effect of high temperature. Consequently, the low-temperature duration may be critical for the immunomodulatory effect of TC-HT. In particular, Fig 5C shows that the HSV-1 clearance efficacy of U-937 macrophages reached its optimum via extension of the low-temperature duration in 42 °C TC-HT to 2 min. Moreover, this optimized 42 °C TC-HT treatment also inhibited the LPS-induced TNF-α and IL-6 and secretions (Fig 6), and enhanced actin polymerization (Fig 7) and phagocytosis activity (Fig 8) in U-937 macrophages, which may lead to the elevated HSV-1 clearance efficacy. Noteworthily, as the low-temperature duration prolongs further, the promoted HSV-1 clearance efficacy gradually declines, and even disappears eventually (Fig 5C). Our previous study also stressed the importance of proper low-temperature duration for therapeutic effect in neuroprotection [31], highlighting its critical role in attaining the optimal therapeutic efficacy.

In summary, this is the first study on the immunomodulatory effect of TC-HT both *ex vivo* and *in vitro*. In the whole blood assay, 42 °C TC-HT post-treatment decreases LPS-stimulated TNF-α and IL-6 levels significantly, which is unseen in 42°C HT post-treatment. Besides, although LPS-stimulated IL-1β level increases after 42 °C TC-HT post-treatment, the extent is significantly milder than that by 42 °C HT post-treatment. Meanwhile, 42 °C TC-HT post-treatment reduces significantly LPS-stimulated TNF-α, IL-6, and IL-1β levels in PBMCs, but 42 °C HT post-treatment shows inhibitive effect only on IL-1β, without such effect on LPS-stimulated TNF-α and IL-6. In addition, 42 °C TC-HT outperforms HT in activating U-937 macrophages to clear HSV-1, and the optimized low-temperature duration in 42 °C TC-HT treatment further boosts the HSV-1 clearance efficacy of U-937 macrophages. Furthermore, it’s noteworthy that TC-HT has greater efficacy in sustaining the cell viabilities of both PBMCs and U-937 macrophages, indicating that TC-HT is a safer and more effective thermal treatment than HT both *ex vivo* and *in vitro*. Overall, the study manifests the immunomodulatory potential of TC-HT on anti-inflammation and viral clearance. Further research on the immunomodulatory mechanism of TC-HT is in the works.

## Acknowledgements

The authors would like to acknowledge the service provided by the Technology Commons, College of Life Science, National Taiwan University for use of flow cytometry system, and express gratitude to Prof. Chun-Jen Chen for using his BSL-2 laboratory for experiments involving HSV-1. We would also like to thank Mr. J. H. Wu for participation of the early phase of this work.

## Author Contributions

**Conceptualization:** Chih-Yu Chao.

**Data Curation:** Yu-Yi Kuo, Chih-Yu Chao.

**Formal analysis:** Yu-Yi Kuo, Guan-Bo Lin, Wei-Ting Chen, You-Ming Chen, Hsu-Hsiang Liu, Chih-Yu Chao.

**Funding acquisition:** Chih-Yu Chao.

**Investigation:** Yu-Yi Kuo, Guan-Bo Lin, Wei-Ting Chen, Chih-Yu Chao.

**Project Administration:** Chih-Yu Chao.

**Supervision:** Chih-Yu Chao.

**Validation:** Yu-Yi Kuo, Guan-Bo Lin, Wei-Ting Chen, You-Ming Chen, Hsu-Hsiang Liu.

**Writing – original draft:** Yu-Yi Kuo, Chih-Yu Chao.

**Writing – review & editing:** Chih-Yu Chao.

